# Chitozyme: First Peroxidase-like Activity of Chitosan for Multiplexed Visual Detection of H_2_O_2_, Glucose and Lactate on Paper-based Device

**DOI:** 10.1101/162446

**Authors:** Syed Rahin Ahmed, Xuan Weng, Suresh Neethirajan

## Abstract

Visual read-out diagnostics tools are promising candidates for field applicable medical devices. Current colorimetric biosensors require introduction of natural enzymes or nanozymes, which has some serious drawbacks for practical applications. Chitosan, a natural polymer, provides safe and efficient compound in medical and pharmaceutical technology. Herein, we report on a simple, cost-efficient, field-portable, environmental friendly and ultra-sensitive multiplex detection platform based on peroxidase-like activity of chitosan in the presence of 3,3’,5,5’-Tetramethylbenzidine (TMBZ) and H_2_O_2_. This straight forward signal amplification strategy was successfully applied to detect H_2_O_2_, glucose and lactate with the limit of detection (LOD) of 2.64 pM, 0.104 μM and 2.8 nM respectively, represents the lowest LOD of H_2_O_2_, glucose and lactate with visual read-out. The chitosan-based assay performance was also retained in complex biological media for glucose and lactate detection. Furthermore, the proposed assay was successfully demonstrated as a paper-based colorimetric biosensor. Most importantly, the simplicity, biocompatibility and sensitivity of the proposed assay will open new doors for instrument free naked eye visual detection of H_2_O_2_, glucose and lactate detection.

## 1. Introduction

Instrument free point of care diagnostic tools has high potential to provide a simple, cheap and easy-accessible device for medical applications. A key example of such device is colorimetric read out strategy that enable the development of low-cost diagnostics to detect analytes visually [1-5]. Over the last few decades, conventional enzymes i.e., horseradish peroxidase (HRP), alkaline phosphate (ALP) have been used for visual color amplification. However, their denaturation behaviour at high temperature, pH stability and high cost limits their use in real life applications.^2^ Recently, researchers are attempting to replace natural enzymes with nanomaterials (known as nanozymes) to overcome the aforementioned problems; and few nanomaterials delivered as promising canditates of this class for various applications [6-8]. Up-to-date, although much efforts have been given on nanomaterials based artificial enzyme developments, their aggregation problems in different chemical environments; monodispersibility; uniform size preparation and toxicological implications are major concerns in the practical applications for in-vitro or in-vivo detection of analytes.

Introducing a new diagnostic tool that is easy-to-operate, containing non-toxic chemicals and ability to detect extremely small analyte concentrations with enhanced superior visual color would enable the development of low-cost commercially viable technologies for medical applications. In view of above discussions, here we propose to amplify the visual detection using chitosan as a peroxidase-like enzyme. We proposed the term of our new approach to be “chitozymes”.

Chitosan has received lots of research interest because of its structural and broader range of chemical and physical attributes in fundamental science, applied research, and industrial biotechnology. For example, chitosan can be used as an advanced biofabrication material; surface adaptation of cell/protein-integrating biological systems; and enzyme surface immobilization in preparation of medical sensing devices [9,10]. Chitosan has benefits of natural availability as well as biocompatibility, brings it as a potential candidate to the creation of chitosan-based analytical tools for point-of-care clinical, biotechnological, and environmental analysis. Recently, the integration of chitosan with nanomaterials was reported as a peroxidase-like activity for visual sensing application [11,12]. However, up-to-now, no reports on bare chitosan as a peroxidase-like enzymatic activity has been published yet. Hence, here for the first time, we present peroxidase-like activity of chitosan for rapid ultrasensitive detection of H_2_O_2_, glucose and lactate. This approach will diminish the requirements of others material to integtrae with chitosan for sensing applications as well as reduce the cost, labour and multi-step preparative process.

Positively charged gold nanoparticles (Au NPs) exhibits higher peroxidase activity in comparison to negatively charged Au NPs [13]. The surface charge of nanoparticles plays a crucial role in influencing the peroxidase-like activity. Hence, we targeted three positively charged chemicals i.e., Cetyl trimethylammonium bromide (CTAB), Poly-L-Lysine (PLL) and chitosan to check their enzymatic activity towards TMBZ/H_2_O_2_. Our results confirm that chitosan demonstrated stronger peroxidase-like activity in comparison to other two chemicals, and allowed us to develop a portable multiplex system for simultaneous detection of glucose, lactate and H_2_O_2_.

Chitosan could catalyse the oxidation of the peroxidase substrate TMBZ by H_2_O_2_ to develop a blue color in aqueous solution, which could introduce us a new way of visual detection of H_2_O_2_. A sensitive and accurate determination of H_2_O_2_ is much needed because of its numerous application in food and pharmaceutical industries, and environmental analysis [14]. In addition, H_2_O_2_ is produced by the oxidation of glucose due to catalysis by glucose oxidase (GOD) and the oxidation of lactic acid catalysed by lactate oxidase (LOD). The proposed sensing mechanism was further extended to quantitatively analyse glucose and lactic acid in samples. The determination of glucose concentration at lower limit is essential for medical diagnostic applications such as for diabetic patients, as well as in industrial applications [15]. The quantitative measurement of lactic acid concentration in body fluids has importance in clinical assessment to determine the patients with diabetic coma, bacterial infections and others medical symptoms [16]. Lactate concentration plays a key parameter in healthcare, food industries and for assessing patient health conditions like hemorrhage, respiratory failure, hepatic disease, sepsis and tissue hypoxia [17].

The creative idea to demonstrate the chitosan-based biosensing assay on a paper-based device towards detecting H_2_O_2_, glucose and lactate would facilitate the pathway for obtaining reliable healthcare data under non-laboratory conditions at low-cost. Recently, paper-based analytical tools has gained attractive alternative to highly sophisticated instrumentation for numerous applications. Paper-based device can be easily printed, coated and impregnated. The cellulose composition in paper is compatible for proteins and biomolecules; it is as well as environmentally compatible, easily disposable and accessible almost everywhere. In particular, paper-based biosensor is highly suitable to point-of-need monitoring in less industrialized countries [18]. Testing of target analytes would simply involve dropping the solutions (blood or serum or sweat or saliva) on chitosan/TMBZ treated paper. This concept can promptly inform even nonprofessional users with clear and unambiguous visual color against target analytes.

## 2. Materials and Methods

### 2.1. Materials

Chitosan, poly-l-lysine (PLL),cetyltrimethylammonium bromide (CTAB), 3,3’,5,5’-tetramethylbenzidine (TMBZ), hydrogen peroxide (H_2_O_2_), sulfuric acid (H_2_SO_4_), glucose, sucrose, fructose, lactose, galactose, glucose oxidase (GOD from Aspergillus Niger), L-Ascorbic acid, L-(+)-Lactic acid, lactic oxidase (LOD from Aerococcus Viridans) and nunc-Immuno 96-well plates were purchased from Sigma-Aldrich (St. Louis, MO, USA). Maltose and acetic acid were received from Fisher Scientific Company (New Jersey, USA). All experiments were performed using highly pure deionized (DI) water (>18 MΩ· cm).

### 2.2 Preparation of water-soluble chitosan

Chitosan solution was prepared by dissolving 2 mg/mL chitosan in 0.1 M glacial acetic acid at 60°C for 30 min. Once the initial turbid colored solution turned to clear one, stopped the heating and cool down at room temperature. Prepared solution was stored at room temperature for further experiments.

### 2.3 Hydrogen peroxide (H_2_O_2_) detection using GOD and chitosan

Various concentrated solutions of H_2_O_2_ were prepared using PBS buffer prior to the sensing experiments. Then, 100 μL of each concentrated H_2_O_2_ solutions were placed in 96 well plate and subsequently, 100 μL of chitosan/TMBZ-H_2_O_2_ mixture solution were added in the above solution. After 30 min, the ultraviolet-visible (UV–vis) spectrum of developed color was recorded using a Cytation 5 spectrophotometer (BioTek Instruments, Inc., Ontario, Canada).

### 2.4 Glucose detection using GOD and chitosan

The experiment for glucose detection was performed as follows: (a) 50 μL of GOD (1 mg mL-^1^) and 50 μL of different concentrated glucose in PBS buffer solution (pH 7.5) were incubated in 96 well plates at room temperature for 30 min; (b) Then, 100 μL of chitosan/TMBZ-H_2_O_2_ mixture solution were added in the above solution, and (c) the mixed solution was incubated at room temperature for 30 min and then the ultraviolet-visible (UV–vis) spectrum of developed color was recorded.

### 2.5 Lactic acid detection using LOD and chitosan

Lactic acid detection was performed as follows: (a) 50 μL of LOD (1 mg mL-1) and 50 μL of different concentrated lactic acid in PBS buffer solution (pH 7.5) were incubated in 96 well plate at room temperature for 30 min; (b) Then, 100 μL of chitosan/TMBZ-H_2_O_2_ mixture solution were added in the above solution, and (c) the mixed solution was incubated at room temperature for 30 min and then the ultraviolet-visible (UV–vis) spectrum of developed color was recorded.

### 2.6 Preparation of paper-based device

Xerox printer (colorQube 8580, Japan) was used to fabricate the paper-based sensing device. Here, Whatman filter paper (GE Healthcare UK limited, Buckinghamshire, UK) was used due to its hydrophilic and biocompatible behavior as well as low cost. The fabrication process includes three steps:

1)The desired geometry i.e., Circle, square and rhombus-like shaped was drawn in white background for H_2_O_2_, glucose and lactic acid detection respectively by using the software Adobe Illustrator.
2)Printed the wax-paper with circle, square and rhombus-like shaped on the surface by using the wax printer.
3)Melted the wax-printed paper on a hot plate at 175°C for 40 s. The wax covered area will be hydrophobic, while the area without wax will be hydrophilic (sensing area).

### 2.7 Detection of H_2_O_2_, glucose and lactic acid on paper-based device

Printed wax-paper was used to detect H^2^O^2^, glucose and lactate. Firstly, 5 μL of chitosan solution (2 mg/mL) was dropped on sensing area (hydrophilic area) for 5 min to dry-up. Then, various concentrated target solutions (5 μL) was added on the surface, and color was developed within 10 min. ImageJ software was used to process the pixel value of the detection analyte from the colored image.

## 3. Results and Discussion

Firstly, the enzymatic activity of chitosan (1 mg/mL, 20 μL), CTAB (1 mg/mL, 20 μL) & PLL (1%, 20 μL) were examined with 1 mL of mixture solution of peroxidase substrate TMBZ (5 mM) and H_2_O_2_ (10 mM) separately for 5 min (Fig. 1A). A strong blue color was developed for chitosan wereas the blue color developed with CTAB was not so significant in comparison to chitosan. No enzymatic change of color was observed for PLL during the reaction time (Fig. 1B). The absorbance peak of resulting blue solution was located at 660 nm due to the oxidation of TMBZ.

**Figure 1.**
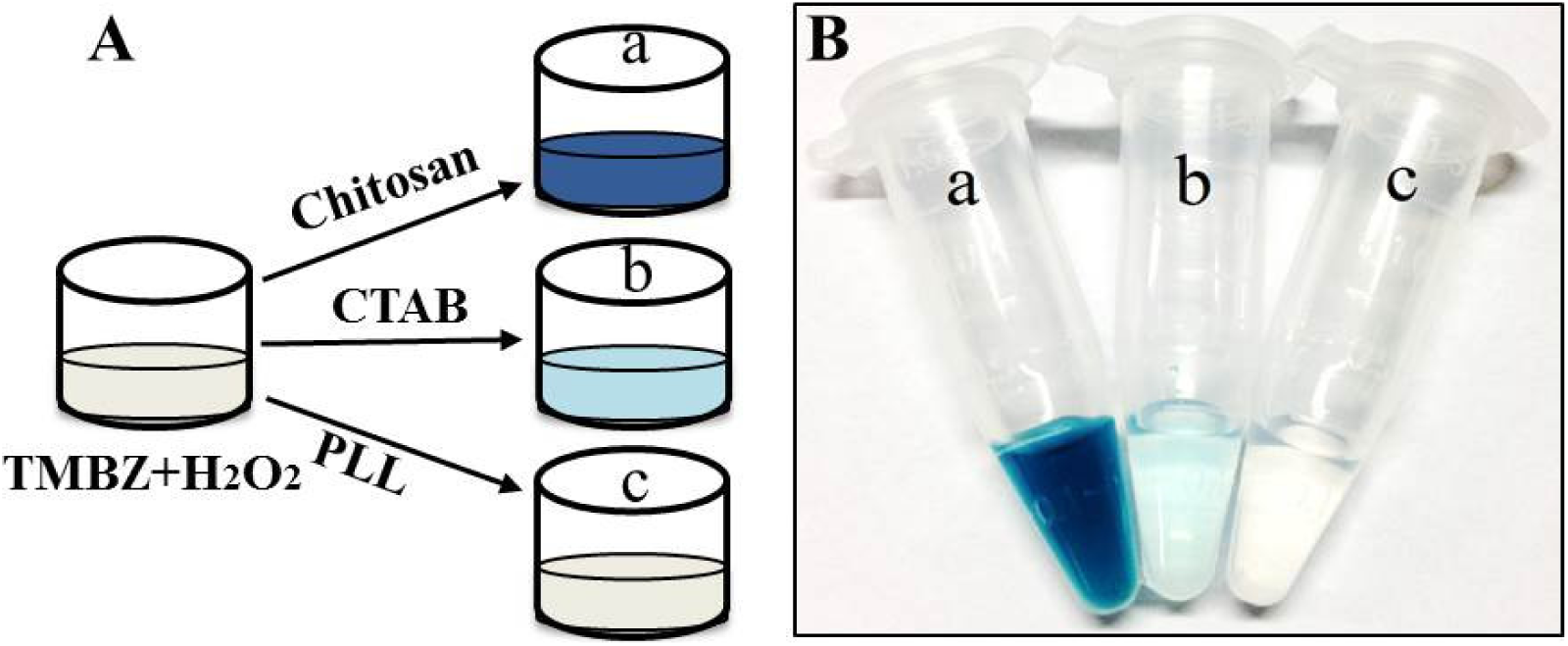
The catalytic activity of chitosan, CTAB and PLL: (A) schematic presentation of the sensing experiment; (B) naked eye image of the experimental results.

The viability of the sensing assay was performed with different kinds of reaction mixtures i.e. a) chitosan/TMBZ- H_2_O_2_; b) chitosan/TMBZ; c) chitosan/H_2_O_2_; d) TMBZ-H_2_O_2_/CH_3_COOH and e) chitosan/CH_3_COOH. As shown in Figure 2A, it was clearly observed with the naked eye that a deep blue colored solution was developed by the reaction of chitosan and TMBZ–H_2_O_2_ complex (Fig. 2A-a). However, none of the other mixtures showed characteristic blue color during reaction (Fig. 2A- b, c,d,e). Here, the viability test of acetic acid (CH_3_COOH) was confirmed since it was used to dissolve chitosan. Spectroscopic study of colored solutions revealed a significant enhanced spectrum peak centered at 660 nm compared to other solutions (Fig. 2B). The control experiments confirmed that the glacial acetic acid did not affect the peroxidase-like activity of chitosan, and it is specific only in the presence of TMBZ-H_2_O_2_ complex.

**Figure 2.**
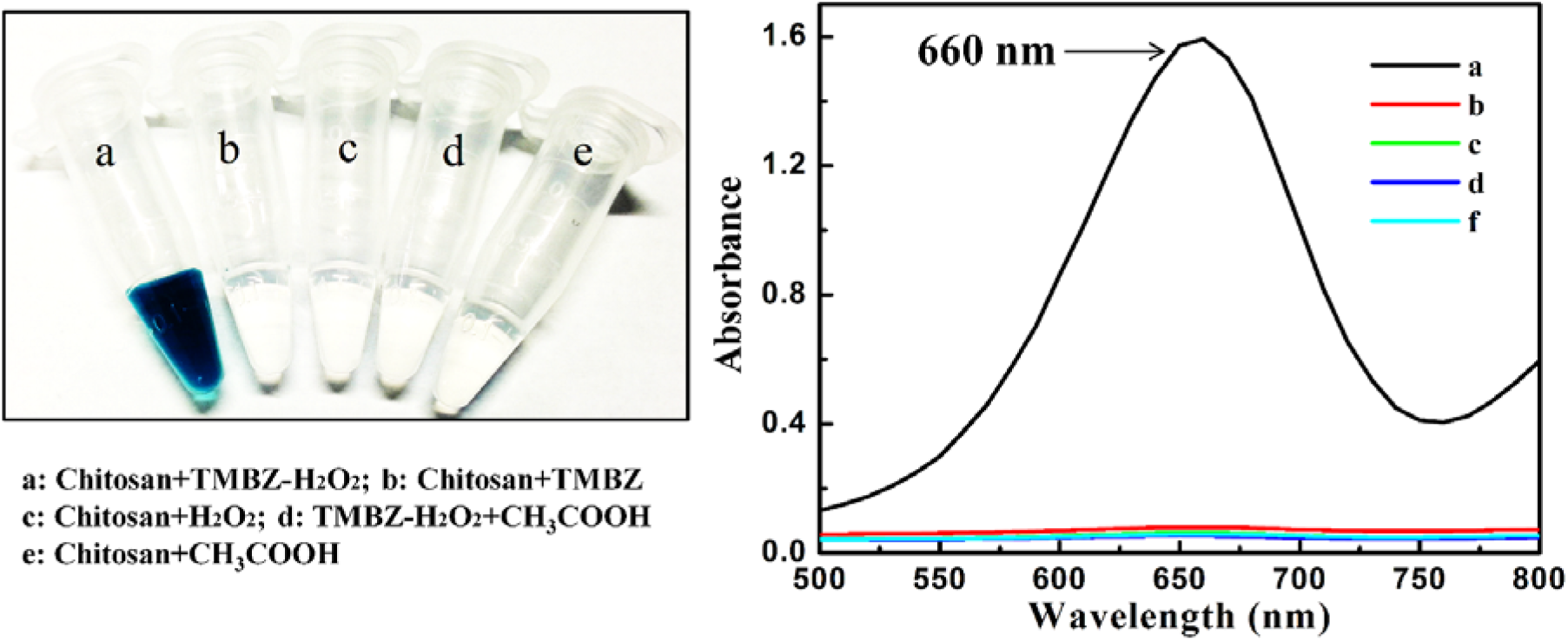
The viability of the developed sensing assay: (A) visual color of the experimental results with different mixtures; (B) Absorbance of the corresponding mixture solutions.

The optimal conditions for catalytic activity of chitosan was examined with several parameters, such as concentration, pH, TMBZ concentration, H_2_O_2_ concentration and reaction time. These parameters may significantly affect the enzymatic activities of chitosan. The concentration of chitosan has significant effect on enzymatic reaction. Experimental studies on different concentrated chitosan showed that the intense blue color was observed with concentration of 2 mg/mL (Fig. S1). The pH effect on the enzymatic activity of chitosan was studied in the range of pH 1–12. Results revealed that the catalytic activity of chitosan increased with increasing the pH up to 4, then sharply decreased with increasing pH values (Fig. S2A). The reaction time to get maximum catalytic activity was investigated from 0 to 20 min (Fig. S2B). The optical density of blue color generated through catalytic activity of chitosan was observed to be a maximum within 9 min, after that, it’s became steady. The optimum concentration of H_2_O_2_ and TMBZ were also checked at 0 to 20 mM, and 1 to 10 mM respectively. The highest peroxidase-like activities of chitosan were observed with the concentration of 10 mM and 5 mM of H_2_O_2_ and TMBZ respectively (Fig. S2C and S2D). All the reactions were carried out at room temperature which is a convenient for sensing applications.

The enzymatic activity of chitosan depends on H_2_O_2_ concentration in the presence of TMBZ. The proposed method was applied to detect H_2_O_2_, and the optical density of experimental results increased with increasing H_2_O_2_ concentration (Fig. 3A). The calibration graph of the optical density at 660 nm versus H_2_O_2_ concentration showed linearity in the range of 10 pM to 1 mM with a detection limit of 2.64 pM (Fig. 3A).

**Figure 3.**
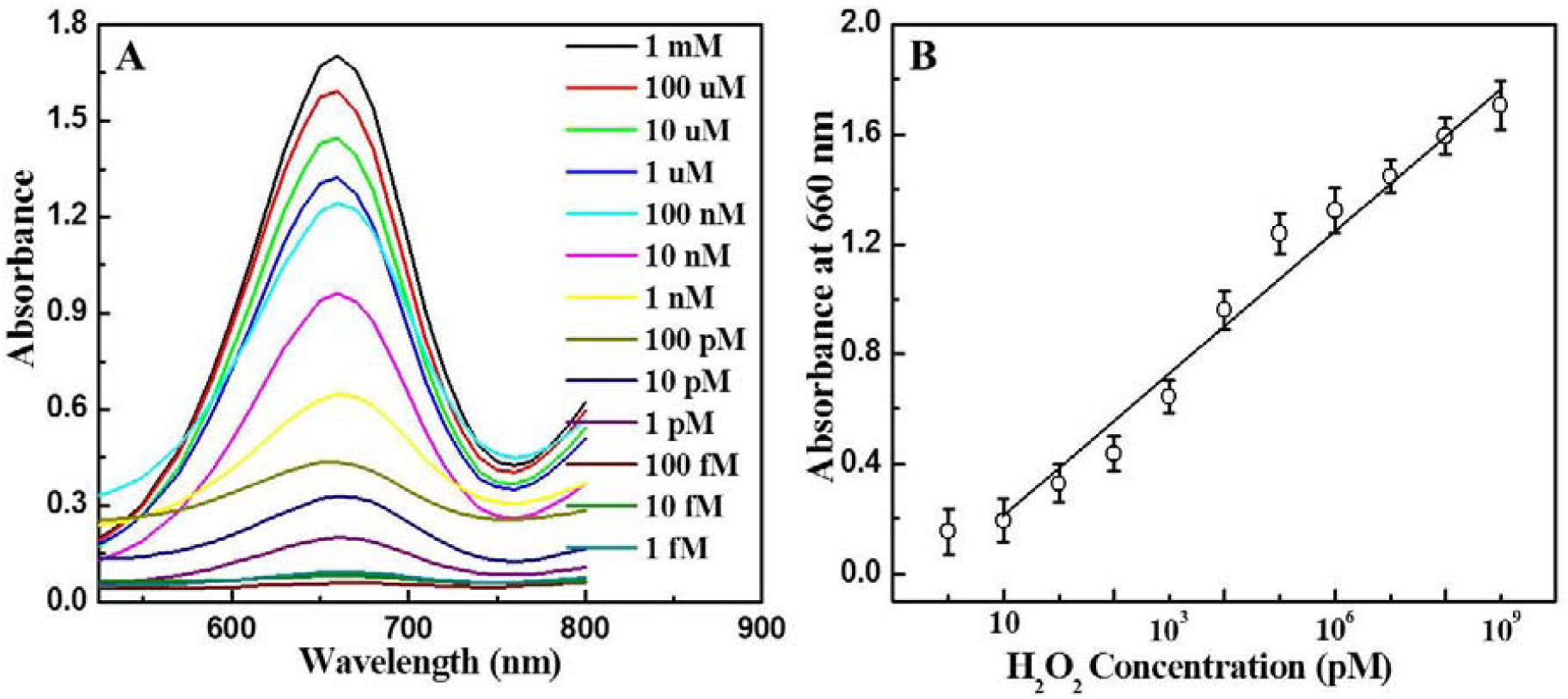
Experimental results of H_2_O_2_ detection: (A) Absorbance spectra of sensing H_2_O_2_; (B) Calibration curve of absorbance Vs H_2_O_2_ concentration.

In addition, the catalytic reaction of chitosan is integrated with the reaction between glucose and GOD for quantitative measurement of glucose concentration (Fig. 4A). The visual detection of glucose was completed by two steps reaction process i.e., at first, glucose and GOD were reacted in PBS (pH 7.4) at room temperature for 10 min which produced H_2_O_2_; then the chitosan was added to detect H_2_O_2_ visually. Fig. 4B showed the linear relationship of the absorbance (660 nm) versus glucose concentration in the range of 0.2-10 μM with the limit of detection of 0.104 μM, which is lower than any paper reported so far to our best of knowledge. The glucose level for diabetic patient when fasting glucose is around 4-7 mM. Thus, our proposed glucose sensor is much sensitive than the practical needs. Several control experiments were also performed to check the specificity of the proposed method for glucose detection using sucrose, fructose, lactose, galactose and maltose. The results were investigated and no characteristic peaks were obtained for control experiments (Fig. S3). Hence, the visual detection method proposed for glucose was highly sensitive and selective. The naked eye image of glucose detection system is shown in Figure S4. Besides the analysis of glucose in phosphate buffer, additional experiments were conducted to ability of detecting glucose in complex biological matrix, such as blood. Results revealed that chitozymes based visual analysis is robust and selective enough to detect glucose in complex media with limit of detection of 0.23 μM, which is promising for practical applications (Fig. S5).

**Figure 4.**
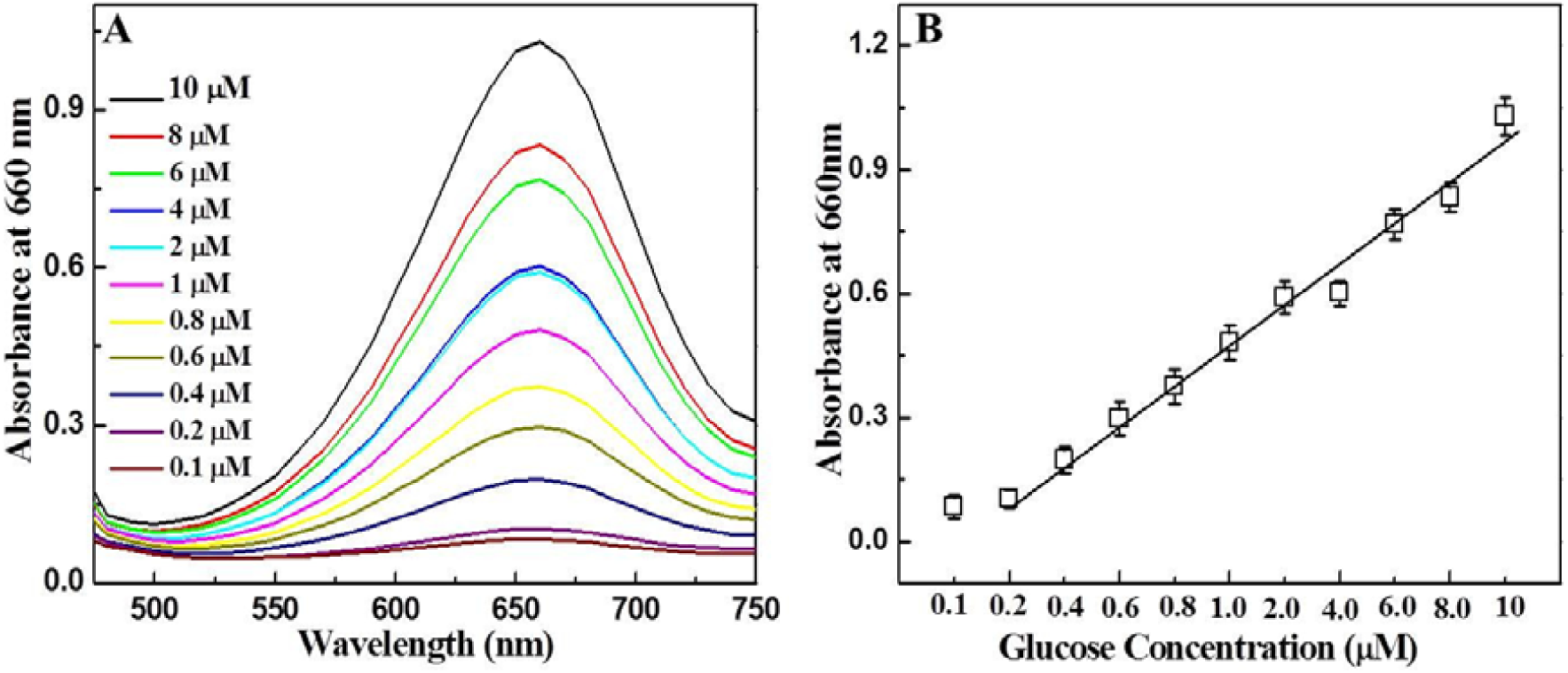
Experimental results of the glucose detection based on the developed Chitozyme biosensing assay: (A) Absorbance spectra of the glucose sensing experiment; (B) Calibration curve of the absorbance Vs glucose concentration.

The analytical reliability of the colorimetric measurements for the glucose assay was compared to the signal recorded in a commercial glucometer (Contour next blood glucose Meter, Japan). The analysis on the glucometer was carried out by the addition of a small drop of target solution in blood media on a disposable test strip. The glucose level appears on the meter display is in mM unit. As shown in Figure S6, commercial glucometer has the capability to detect glucose concentration with the range of 4-10 mM; whereas the proposed method showed its sensitivity up to μM level.

Furthermore, similar two step assay was conducted for lactic acid detection using LOD to produce H_2_O_2_ at the initial step (Fig. 5A). The absorbance of sensing results was linearly correlated with lactic acid in the range of 10 mM to 10 nM with the limit of detection of 2.8 nM (Fig. 5B). While lactic acid analysis was performed with commercial milk, detection limit was obtained at 5.3 nM (Fig. S7). Thus, the developed chitozymes based analysis of lactic acid would be a promising candidate for practical applications in complex media. A comparison study with others techniques was also performed based on recently published papers as shown in Table S1 & S2. Superiority in terms of sensitivity of the present study in comparison to other techniques was clearly observed here.

**Figure 5.**
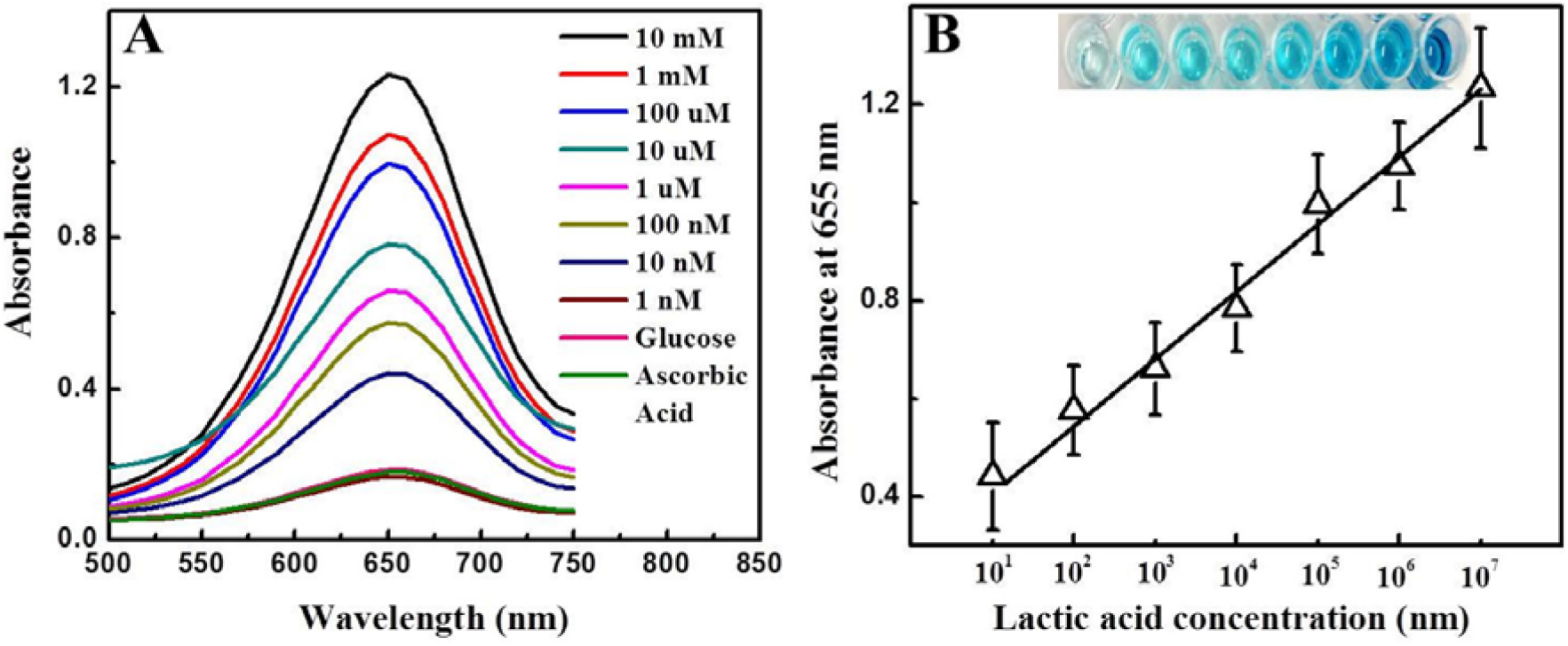
Experimental results of the lactic acid detection based on the developed chitozyme biosensing assay: (A) Absorbance spectra of the lactic acid sensing (Inset: visual detection image); (B) Calibration curve of the absorbance vs lactic acid concentration.

Proposed chitozymes based colorimetric detection method was further demonstrated with paper-based sensor to make it more flexible in real life applications. Paper based disposable devices has been investigated extensively recently due to low-cost and field-portable in remote area applications [19]. Here, paper device was prepared by wax printer with desired geometry. Circle, square and rhombus-like shaped (diameter of 8 mm each) was drawn and printed for H_2_O_2_, glucose and lactic acid detection respectively. The bottom part of the paper devices was covered with wax to prevent sample leaking.

The desired detection zone on printed wax paper device was initially modified with chitosan solution (5 μL) and allowed to dry at room temperature for 10 min. Then, the detection zone was spotted with different concentrated H_2_O_2_, mixture containing different concentrated glucose and GOx; mixture containing different concentrated lactic acid and LOD prepared in PBS buffer solution (pH 7.5) separately. Visual image of detections on paper-based device is shown in Figure 6A. The linear concentration range for H_2_O_2_ detection was 1 mM to 100 nM with the limit of detection of 64.3 nM (Fig. 6B); for glucose detection was10 μM to 0.6 μM with the limit of detection of 0.46 μM (Fig. 6C) and for lactic acid, the detection was10 mM to 10 nM with limit of detection 7.9 nM (Fig. 6D).

**Figure 6.**
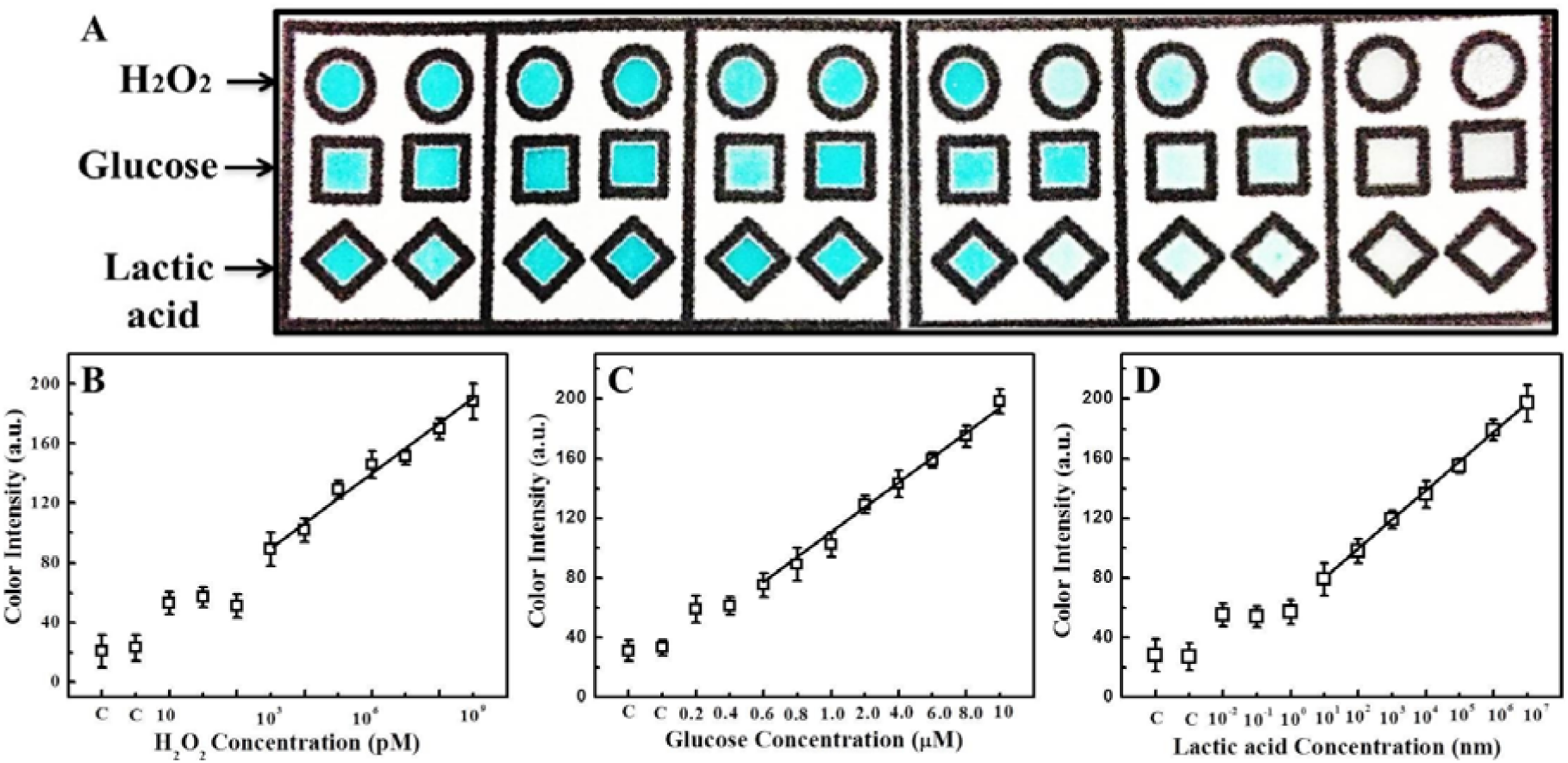
Experimental results of the chitozyme based paper-based biosensor: (A) Visual detection of H_2_O_2_, glucose and lactic acid; (B) Calibration curve of color intensity vs H_2_O_2_ concentration (C means control experiment with ascorbic acid & glucose); (C) Calibration curve of color intensity vs glucose concentration (C means control experiments with fructose & sucrose); (D) Calibration curve of color intensity vs lactic acid concentration (C means control experiments with ascorbic acid & glucose)

## 4. Conclusions

In conclusion, an user and environment friendly colorimetric read-out assay was introduced based on peroxidase-like enzymatic activity of chitosan for the first time. This straight forward signal amplification strategy was successfully applied to detect H_2_O_2_, glucose and lactic acid with the limit of detection of 2.64 pM, 0.104 μM and 2.8 nM respectively which was the lowest value so far reported to the best of our knowledge. Proposed assay showed its superiority and practicability in detecting the target analytes in the complex media. Chitosan-based bioassay was successfully demonstrated on the paper-based device to make a cheaper and easier medical tool. Because chitosan is a natural and biocompatible compound, this approach would open a new window in colorimetric bioassay and catalytic chemistry. The near future will be an exciting period in the chitosan-based visual biosensor research area.

## Supplementary Materials

The following are available online at http://www.mdpi.com/link, Figure S1-S7, Table S1 and Table S2.

## Acknowledgments

The authors sincerely thank the Natural Sciences and Engineering Research Council of Canada (400705) for funding this study.

## Author Contributions

SN conceived the study. SRA conducted experiments and collected data. XW helped with the fabrication of paper based biosensing device. SRA wrote the manuscript with XW. All authors read and approved the manuscript.

